# Fibroblast-encoded inflammatory memory orchestrates recurrent skin inflammation *via* NNMT-dependent metabolic remodeling

**DOI:** 10.64898/2026.04.27.721223

**Authors:** Yifan Zhou, Kang Li, Yuming Xie, Chencan Su, Zhiguo Li, Yang Yang, Pei Qiao, Wanting Liu, Yaxing Bai, Ke Xue, Johann E. Gudjonsson, Gang Wang, Shuai Shao

**Author notes:** These authors contributed equally to this study (Yifan Zhou, Kang Li, Yuming Xie, Chencan Su). Correspondence: Shuai Shao, MD, PhD, Department of Dermatology, Xijing Hospital, Fourth Military Medical University, 127 Changlexi Road, Xi’an 710032, China.; Or Gang Wang, MD, PhD, Department of Dermatology, Xijing Hospital, Fourth Military Medical University, 127 Changlexi Road, Xi’an 710032, China.; Or Johann E. Gudjonsson, MD, PhD, Department of Dermatology, University of Michigan, Ann Arbor, MI, 48109, USA.

## Abstract

Chronic skin inflammation frequently recurs at the same anatomical sites after therapy withdrawal, implying stromal cells may encode local inflammatory memory. Here, we identified nicotinamide N-methyltransferase (NNMT) as a central metabolic-epigenetic regulator of fibroblast inflammatory memory enabling psoriasis relapse. Single-cell and spatial transcriptomics revealed that dermal fibroblasts acquire a persistent senescence-associated secretory phenotype (SASP) during inflammation, maintaining pro-inflammatory niche in resolved skin that supports CD 8^+^CD 103^+^ tissue-resident memory T cell (Trm) differentiation. Multi-omics profiling demonstrated that NNMT depletes S-adenosylmethionine (SAM), reduces H 3K 27me3 deposition, and permits sustained AP-1 occupancy at SASP gene promoters. Fibroblast-specific NNMT ablation or pharmacologic inhibition suppressed SASP activity, limited Trm accumulation, and prevented both initiation and relapse of skin inflammation in mice. These findings establish NNMT as a stromal regulator linking fibroblast metabolism to durable epigenetic memory and propose its targeting to erase inflammatory memory and achieve long-term remission in psoriasis and related immune-mediated diseases.

Graphic Abstract:
Proposed mechanism through which NNMT-driven fibroblast SASP facilitates the initiation and recurrence of psoriasis

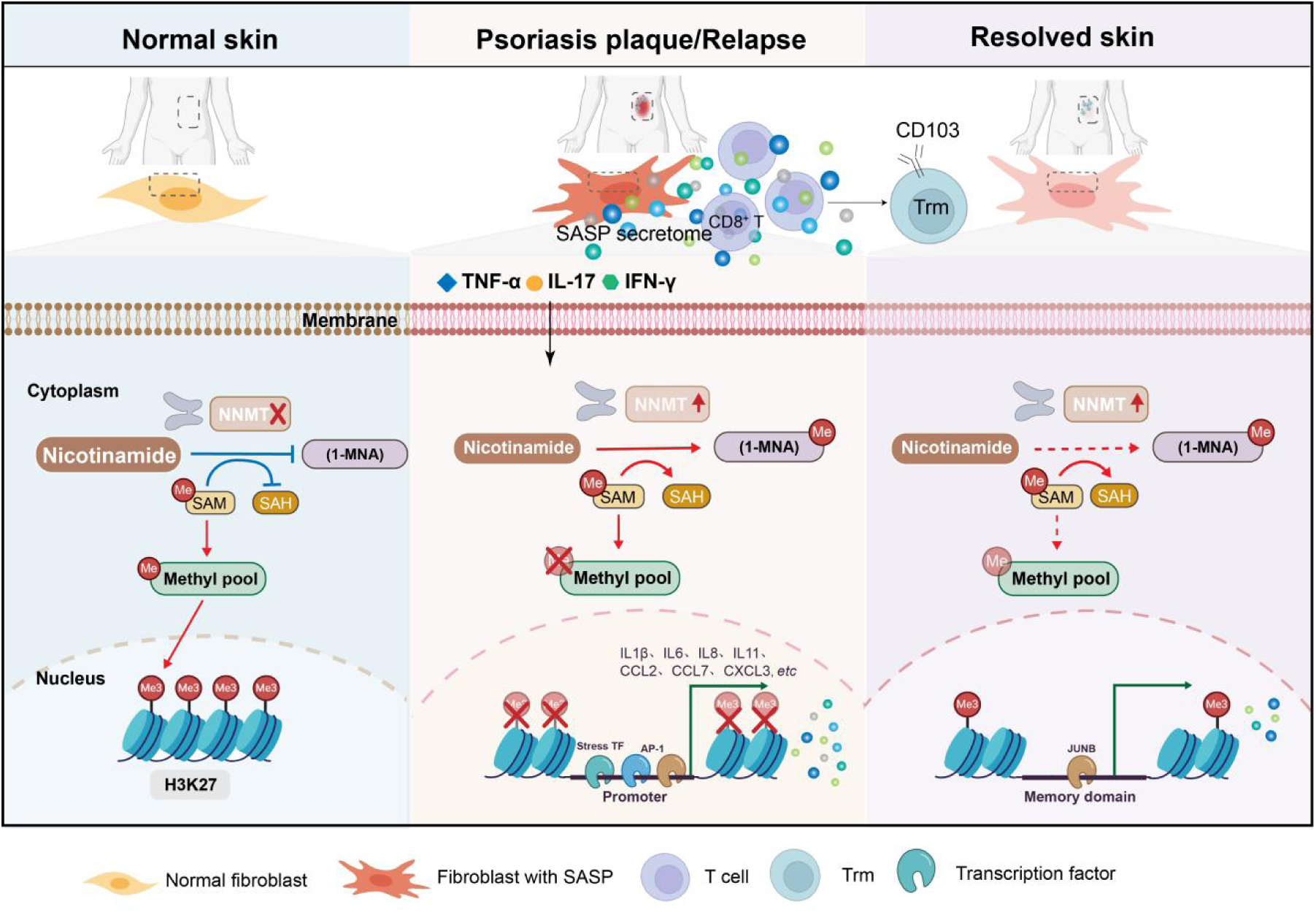

## Introduction

Immune-mediated skin conditions, such as psoriasis, are characterized by recurrent flares that persist despite cytokine blockade therapy^1^. Although inhibition of IL-17 or IL-23 can induce remission, relapse often occurs within weeks to months after treatment withdrawal, especially at the same anatomical sites^2, 3^. This site-specific recurrence implies that cutaneous tissues harbor a form of local inflammatory memory independent of ongoing immune activation^4^. Recent studies have highlighted the contribution of tissue-resident immune cells to this phenomenon. CD 8⁺CD 103⁺ tissue-resident memory T cells (Trm) persist in clinically resolved psoriatic skin and can rapidly produce IL-17 and IL-22 upon reactivation, driving keratinocyte hyperproliferation and lesion recurrence^5, 6^. Similarly, CCR 7⁺ dendritic cells^7^ and γδT17 cells^8^ maintain low-level inflammatory readiness even after apparent resolution. However, biologic blockade of these immune pathways, while highly effective, rarely provides durable response, suggesting that non-immune stromal cells may retain inflammatory memory capable of reigniting local inflammation.

Among these structural cells, dermal fibroblasts are emerging as pivotal regulators of tissue inflammation. They orchestrate immune cell recruitment and cytokine production and can adopt persistent inflammatory states in chronic disease. Our previous work and others have identified fibroblast subsets (*e.g.*, COL6A5⁺ WNT5A⁺ fibroblasts) that amplify psoriasis-associated immune circuits^9, 10^. In other tissues, fibroblasts exhibit features of cellular senescence and a senescence-associated secretory phenotype (SASP), secreting pro-inflammatory cytokines, chemokines, and matrix-remodeling enzymes that perpetuate chronic inflammation^11, 12^. Whether dermal fibroblasts in psoriasis acquire similar long-term SASP features and whether such changes contribute to inflammatory memory remains unknown. At the mechanistic level, epigenetic regulation is central to cellular memory formation. Persistent histone modifications, such as H 3K 27me3 or H 3K 4me1, can prime chromatin for rapid transcriptional reactivation^13^. Inflammatory stimuli can therefore “train” structural cells, endowing them with a memory-like phenotype even after resolution^14^. However, how this epigenetic memory is metabolically sustained is unclear.

Here, we show that dermal fibroblasts in inflamed skin acquire a durable SASP that persists in clinically resolved tissue and functions as a stromal reservoir of inflammatory memory. Using integrated multi-omics profiling, including RNA-seq, Assay for Transposase-Accessible Chromatin with high throughput sequencing (ATAC-seq), and Cleavage Under Targets and Tagmentation (CUT&Tag), and *in vivo* genetic models, we identified Nicotinamide N-methyltransferase (NNMT) as a central regulator of the fibroblast SASP program. By consuming S-adenosylmethionine (SAM), NNMT reduces H 3K 27me3 deposition and maintains chromatin accessibility at SASP gene promoters, enabling persistent AP-1 (*e.g.* JunB)-mediated transcriptional activation. Fibroblast-specific deletion or pharmacologic inhibition of NNMT effectively erases this stromal memory, suppresses accumulation of CD 8^+^CD 103^+^ Trms, and prevents local relapse of skin inflammation. These findings redefine fibroblasts as active organizers of tissue inflammatory memory and establish NNMT as a therapeutic target for achieving durable remission in skin inflammation.

## Results

### Fibroblast SASP persists as stromal inflammatory memory in psoriatic and resolved skin

To determine whether dermal fibroblasts contribute to inflammatory memory in psoriasis, we first analyzed single-cell RNA sequencing (scRNA-seq) datasets from lesional, non-lesional, resolved, and relapsed skin of psoriasis patients. Using the SenMayo gene set^15^ as a reference, both AddModuleScore and AUCell algorithms consistently identified fibroblasts as the dominant SASP-scoring population, followed by myeloid and endothelial cells (Fig. S1a-b, Fig. 1a, and Supplementary Table 1). Within the fibroblast compartment, SASP scores were markedly elevated in active psoriatic lesions compared with healthy controls and non-lesional skin (Fig. 1b and Fig. S1c). Notably, fibroblasts isolated from clinically resolved lesions after 40 weeks of successful anti-IL-17A therapy retained significantly higher SASP scores than those from healthy skin, and fibroblasts from relapsed lesions exhibited the highest levels overall (Fig. 1b). These findings indicate that the SASP program persists in clinically resolved psoriatic skin, suggesting fibroblast-intrinsic inflammatory memory.

**Figure 1.**
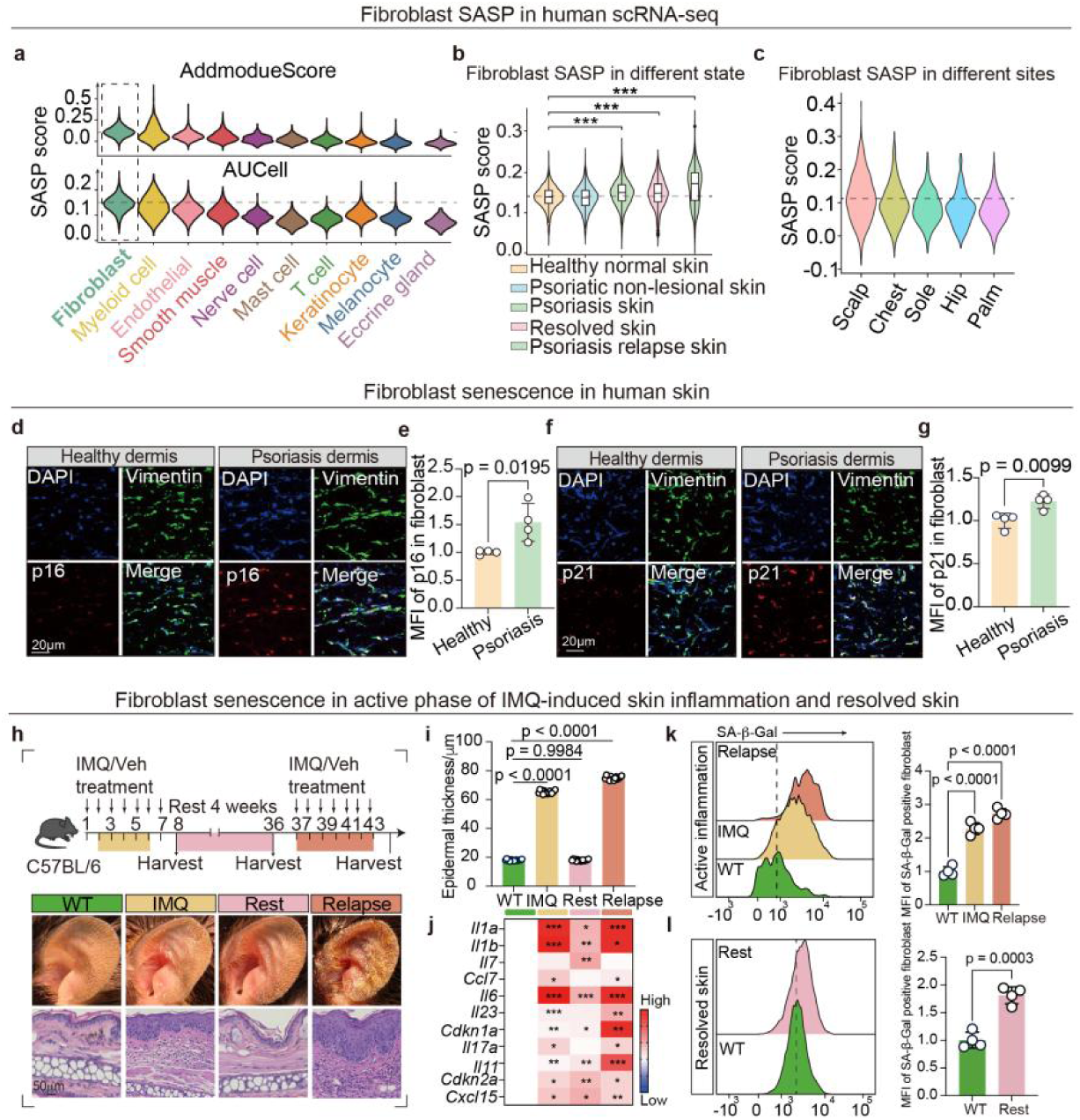
Fibroblast SASP persists as stromal inflammatory memory. **(a-c)** Single-cell analysis showing SASP scores across skin cell types (a), fibroblast SASP scores in healthy normal skin, psoriatic non-lesional skin, psoriatic skin, resolved skin, and psoriasis relapse skin (b), and fibroblast SASP scores in different body sites (c). **(d-g)** Representative tissue immunofluorescence and MFI quantification of p16 (d-e) and p21 (f-g) in skin fibroblasts from healthy individuals and psoriasis patients (n= 4 skin samples/group). **(h)** Schematic of IMQ-induced psoriasis, rest, and relapse mouse models (the upper panel) and representative macroscopic images and H&E images (the lower panel). **(i)** Statistical analysis of epidermal thickness (n= 12 of 4 skin samples/group). **(j)** Heatmap showing the expression of inflammatory genes by qRT-PCR during the psoriasis phase, rest phase, and relapse phase, compared with the normalized expression in wild type (n= 4 skin samples/group). **(k-l)** Representative images and MFI quantification of flow cytometry showing increased SA-β-Gal activity in fibroblasts from IMQ and relapse models (k), and SA-β-Gal in fibroblasts from WT mice and IMQ mice after four weeks of remission (l) (n= 4 skin samples/group). SASP, senescence-associated secretory phenotype. MFI, mean fluorescence intensity. IMQ, imiquimod. Veh, vehicle. SA-β-Gal, senescence-associated β-galactosidase. Data are represented as mean ± SD. Analysis of data in (b), (i), and (k) was analyzed using one-way ANOVA with Tukey’s post hoc test. Data in (e), (g), and (l) were analyzed using an unpaired, two-tailed Student’s t-test. *p< 0.05, **p< 0.01, ***p< 0.001.

Spatially, fibroblast SASP activity was enriched in difficult-to-treat sites such as the scalp (Fig. 1c), consistent with areas prone to relapse. Across multiple inflammatory and fibrotic skin disorders, including scleroderma, lupus, and keloids, fibroblast SASP scores were similarly elevated (Fig. S1d), indicating that this state represents a shared stromal program across chronic inflammatory skin diseases.

To validate these observations, immunofluorescence staining confirmed increased expression of senescence markers p16^INK 4A^ (coded by CDKN 2A) and p21 (coded by CDKN 1A) in fibroblasts within psoriatic dermis compared with healthy controls (Fig. 1d-g and Fig. S1e-f). To determine whether this phenotype persists *in vivo*, we established a three-phase imiquimod (IMQ) mouse model comprising induction, remission, and relapse (Fig. 1h). Consistent with the human data, there are obvious scales on the ears of the mice, and the thickness of the ears has significantly increased in the groups of IMQ and relapse (Fig. 1h-i). Quantitative real-time polymerase chain reaction (qRT-PCR) of skin tissue revealed elevated expression of inflammatory cytokines (*Il17a*, *Il23,* and *Il6, etc.*) (Fig. 1j), and the mean fluorescence intensity (MFI) of SA-β-Gal in fibroblasts also significantly increased during the active inflammation phase (Fig. 1k). However, although macroscopic inflammation resolved after four weeks of rest (Fig. 1h-i), fibroblasts from resolved skin continued to express high levels of critical pro-inflammatory cytokines, including *Il6*, *Il1b*, *Il7,* and the senescence markers *Cdkn1a* and *Cdkn2a* (Fig. 1j), and exhibited persistent SA-β-Gal activity (Fig. 1l). These results demonstrate that fibroblasts acquire a SASP phenotype during inflammation that persists after resolution, providing a stromal memory reservoir that may predispose to relapse.

### SASP Fibroblasts establish a pro-inflammatory niche that promotes Trm differentiation and relapse

We next examined how SASP fibroblasts shape the immune microenvironment during disease progression. We divided 11,152 fibroblasts across disease states into three groups based on the SASP score: senescent (top 40%), stable (middle 20%), and non-senescent subsets (bottom 40%). Pseudotime trajectory analysis indicated that senescent fibroblasts emerged early in non-lesional skin and expanded dominantly in psoriatic lesions (Fig. 2a), suggesting progressive acquisition of the SASP phenotype. CellChat network inference showed that senescent fibroblasts exhibited markedly increased outgoing signaling strength, positioning them as central signaling hubs within psoriatic tissue (Fig. 2b and Fig. S2a). GO enrichment analyses of deferentially expressed genes (DEGs) between senescent *vs.* non-senescent fibroblasts revealed enrichment in pathways related to Th1/Th17 responses, leukocyte recruitment, and cytokine secretion (Fig. 2c and Fig. S2b), consistent with their capacity to sustain chronic inflammation.

**Figure 2.**
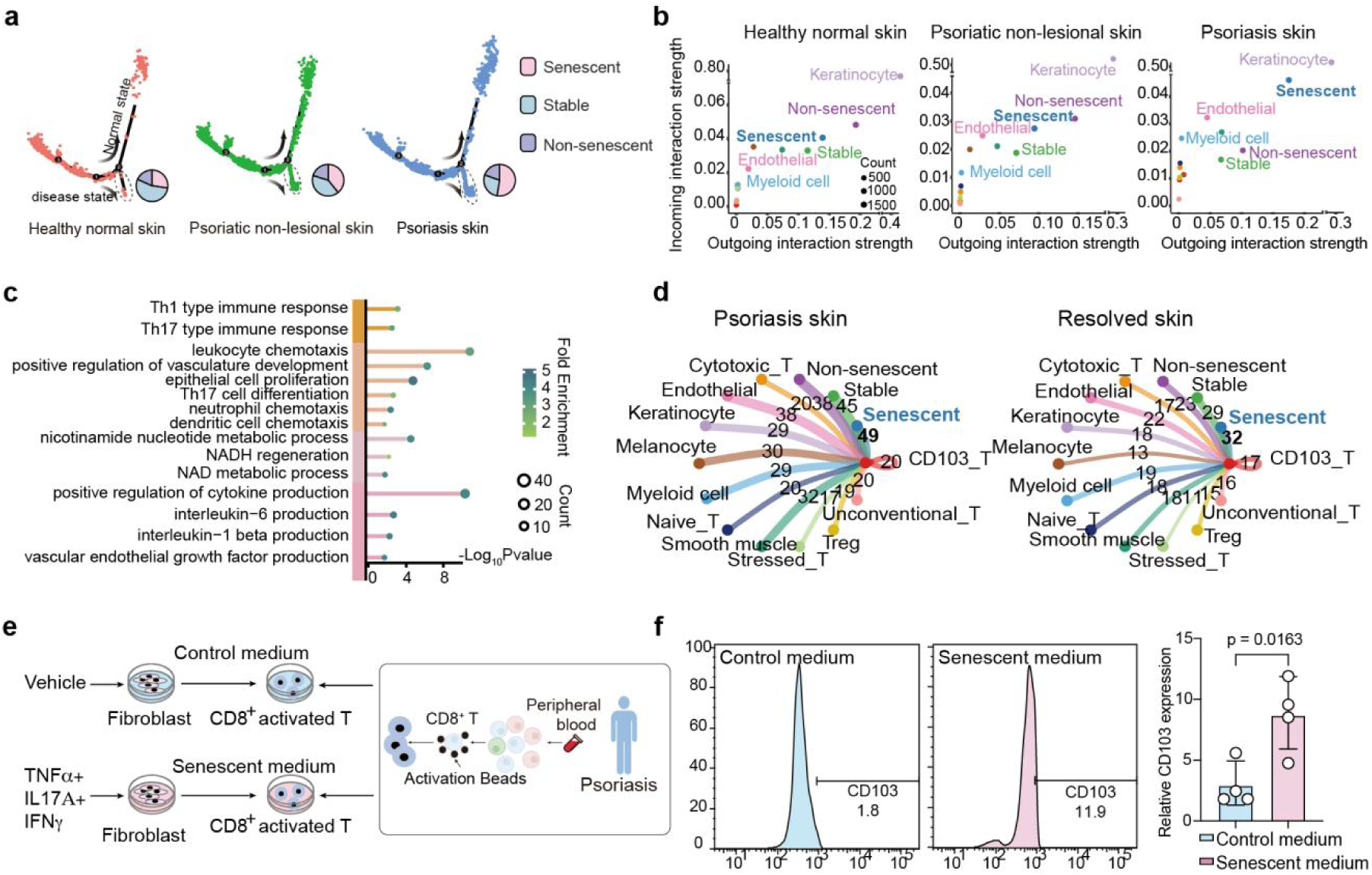
SASP fibroblasts remodel the immune niche to promote Trm persistence. **(a)** Pseudotime trajectory of fibroblasts from healthy skin, psoriatic non-lesional skin, and psoriatic skin; pie charts show proportions of senescent, stable, and non-senescent subpopulations along the branch. The psoriatic branch is circled by a dashed line. **(b)** Scatter plots of intercellular communication networks across the three skin states: x-axis, outgoing signaling; y-axis, incoming signaling; dot size reflects connection numbers, and colors distinguish cell groups. **(c)** GO enrichment of DEGs in senescent versus non-senescent fibroblasts (bubble size, gene count; color, fold enrichment). **(d)** Communication analysis of CD 103⁺ Trm as receiver cells in psoriatic and resolved skin (40 weeks post–anti-IL-17A), showing absolute numbers of incoming signaling pathways from each cell type. **(e)** Schematic of establishing *in vitro* SASP models and coculturing with activated CD 8⁺ T cells from psoriasis patients. **(f)** Representative flow cytometry plots and quantification of CD 103 expression on CD 8⁺ T cells stimulated with control or senescent-fibroblast supernatant (n= 4 blood samples/group). Senescent, stable, and non-senescent denote fibroblast subgroups. Data are represented as mean ± SD. Analysis was performed using an unpaired, two-tailed Student’s t-test.

Given the well-established role of CD 8^+^CD 103^+^ Trm cells in psoriatic recurrence^6^, we assessed fibroblast-Trm communication. CellChat analysis identified senescent fibroblasts as the dominant sender population targeting Trm cells in both inflamed and clinically resolved skin following IL-17A blockade for 40 weeks (Fig. 2d). Similar fibroblast-Trm interactions persisted even after 14 days of anti-IL-23 therapy (Fig. S2c), suggesting that stromal-immune crosstalk remains active despite clinical improvement.

To functionally test this interaction, we established a human dermal fibroblast SASP model *in vitro*. Gene Set Enrichment Analysis (GSEA) identified TNF-α, IL-17A, and IFN-γ as upstream cytokines driving SASP induction (Fig. S2d). Fibroblasts treated with combined-cytokines displayed elevated SA-β-Gal activity (Fig. S2e) and increased expression of inflammatory mediators, including *IL1B*, *IL6*, *IL7*, *CXCL8*, *IL11* and *CCL2* (Fig. S2f), confirming SASP induction. Conditioned media from SASP fibroblasts induced robust CD 103 expression on pre-activated CD 8⁺ T cells isolated from psoriasis patients (Fig. 2e-f and Fig. S2g), indicative of Trm differentiation. Together, these findings demonstrate that senescent fibroblasts actively remodel the immune landscape, establishing a SASP niche that promotes Trm differentiation and persistence, thereby predisposing skin to relapse.

### NNMT acts as the central driver of fibroblast SASP

To identify upstream regulators controlling fibroblast SASP, we compared transcriptional profiles between senescent and non-senescent fibroblasts. Among all DEGs, nicotinamide N-methyltransferase (NNMT) emerged as the most significantly upregulated gene (Fig. 3a). Its expression was largely confined to fibroblasts, with minimal expression in endothelial cells or smooth muscle cells (Fig. 3b). Within the fibroblast compartment, NNMT expression correlated tightly with SASP scores, peaking in the senescent subset in psoriatic skin (Fig. 3b). Across inflammatory and fibrotic disorders including scleroderma, keloids and normal scar, NNMT expression was similarly elevated (Fig. S3a), indicating a shared stromal activation program. Spatial transcriptomics (Fig. 3c) and tissue immunofluorescence (Fig. 3d-e) confirmed expansion of NNMT^hi^ fibroblasts in both papillary and reticular dermis of inflamed psoriatic skin (Fig. S3b and Supplementary Table 2).

**Figure 3.**
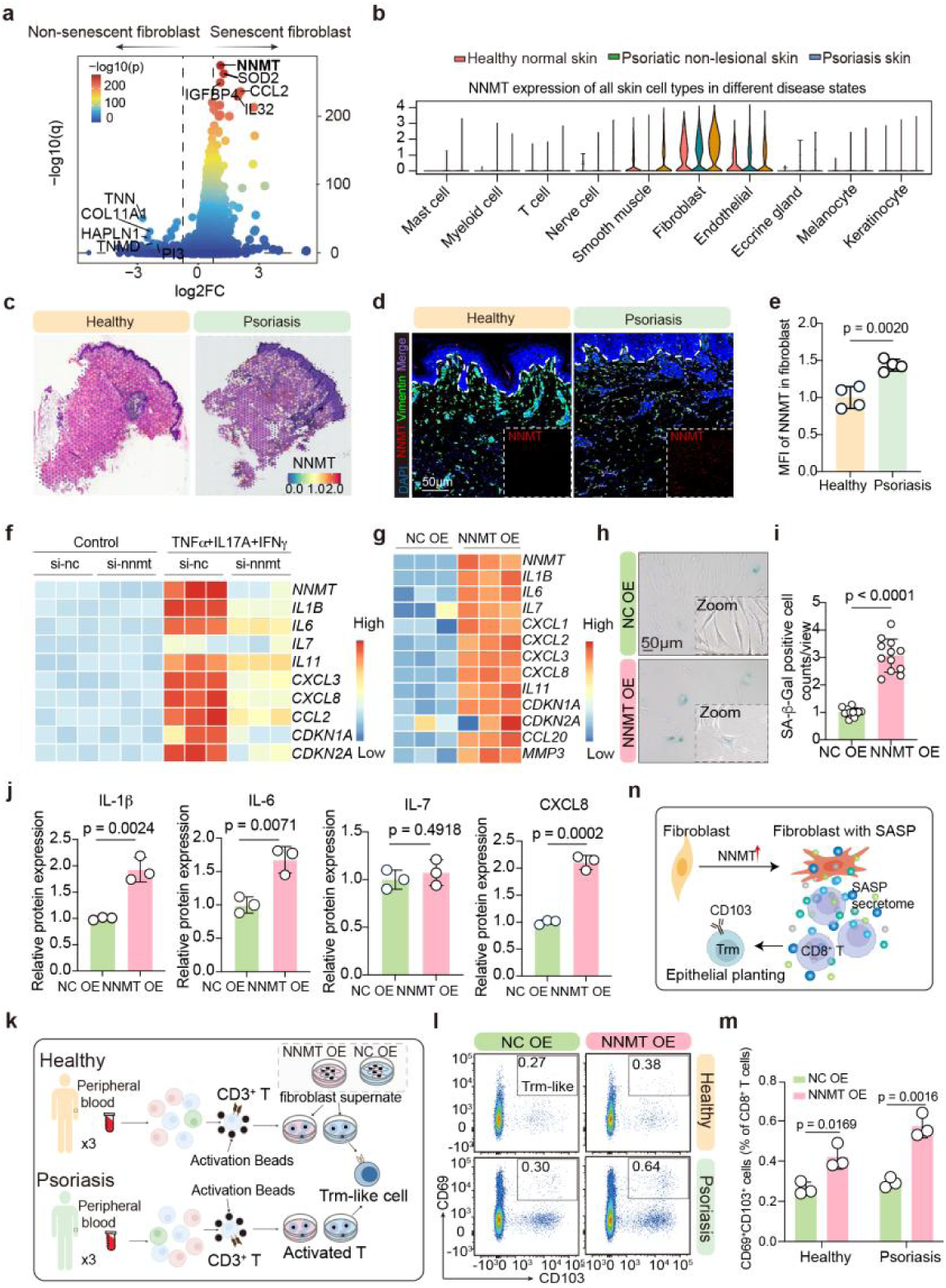
NNMT is a central regulator of fibroblast SASP. **(a)** Volcano plot of DEGs between senescent and non-senescent fibroblast subpopulations (BH-adjusted p/q values; FC, fold change). **(b)** Violin plot of NNMT expression across skin cell states. **(c)** Spatial transcriptomic map of NNMT expression. **(d-e)** Immunofluorescence staining and quantification of NNMT in healthy versus psoriatic skin (n= 4 skin samples/group). **(f-g)** Heatmaps of SASP gene expression by qRT-PCR after NNMT knockdown (f) or overexpression (g) in fibroblasts (n= 3 cell samples/group). **(h-i)** Representative images and quantification of SA-β-Gal staining in NNMT OE versus NC OE fibroblasts (n= 12 views from 4 cell samples/group). **(j)** ELISA of IL-1β, IL-6, IL-7, and CXCL8 in supernatants from NC OE and NNMT OE fibroblasts (n=3 cell samples/group). **(k)** Schematics of inducing CD 8^+^CD 69^+^CD 103^+^ Trm-like cells *in vitro.* **(l-m)** Flow-cytometry plots (l) and quantification (m) of Trm-like cell proportions in peripheral blood T cells from healthy individuals and psoriasis patients treated with NC/NNMT OE fibroblast supernatants (n= 3 blood samples/group). **(n)** Schematic model of NNMT-driven fibroblast SASP promoting Trm differentiation. NNMT, nicotinamide N-methyltransferase; OE, overexpression; NC, negative control; Trm, tissue-resident memory T cells. Data are represented as mean ± SD. Data in (e), (i), and (j) were analyzed using an unpaired, two-tailed Student’s t-test. Analysis of data in (m) was performed using two-way ANOVA.

To functionally validate NNMT’s contribution to the SASP phenotype, siRNA-mediated knockdown of NNMT (Fig. S3c-d) was used in cytokine-stimulated fibroblasts SASP model and showed markedly reduced SA-β-Gal activity (Fig. S3e-f) and mRNA expression of SASP mediators including *IL1B*, *IL6*, *CXCL3*, *IL11*, *CXCL8* and *CCL2* (Fig. 3f). Conversely, NNMT overexpression (Fig. S3g-h) induced enhanced SASP cytokine mRNA expression (Fig. 3g and Supplementary Table 3), elevated SA-β-Gal staining level with cellular enlargement, more indistinct boundaries (Fig. 3h), increased p16^INK 4A^/p21 expression (Fig. S3i-k), and secretion of IL-1β, IL-6, IL-7, and CXCL8 (Fig. 3j). Importantly, conditioned media from Notably, NNMT-overexpressing fibroblasts promoted the differentiation of CD 8⁺CD 69⁺CD 103⁺ Trm-like cells *in vitro* from both healthy and psoriatic donors (Fig. 3k-m and Fig. S3l). These data identify NNMT as the master regulator of fibroblast senescence and SASP, linking metabolic reprogramming to immune activation (Fig. 3n).

### Fibroblast-specific NNMT deletion attenuates initiation of inflammatory responses in skin and prevents relapse *in vivo*

To directly assess NNMT’s functional relevance *in vivo*, we generated a fibroblast-specific NNMT knockout model using an Adeno-Associated Virus expressing short hairpin RNA targeting Nnmt (AAV-shNnmt) delivery under the *Col1a2*-Cre^ERT^ (Fig. 4a-b). Effective gene silencing was confirmed by immunostaining (Fig. S4a) and flow cytometry (Fig. S4b-c). Following IMQ challenge, fibroblast-specific Nnmt knockdown (Nnmt^fibKD^) mice exhibited markedly attenuated erythema and scaling compared with negative control knockdown (NC^fibKD^) mice (Fig. 4c), accompanied by reduced ear thickness (Fig. 4d), epidermal thickness (Fig. 4e), infiltrated immune cells (Fig. 4f), and inflammatory cytokine expression (Fig. 4g). Flow cytometric analysis demonstrated a pronounced reduction in the differentiation (Fig. 4h-i and Fig. S4d) and maintenance (Fig. S5a) of CD 8⁺CD 103⁺ Trm cells in Nnmt^fibKD^ mice relative to controls during “IMQ-Rest-Relapse” phases. The spleen CD 44^+^ memory T cells proportion also significantly decreased in Nnmt^fibKD^ mice (Fig. S5b-c), both for CD 4^+^ (Fig. S5d-e) and CD 8^+^ memory T cells (Fig. S5d, f), and this effect was more pronounced during the remission period. Besides, the reduction of systemic inflammation in the Nnmt^fibKD^ mice was also reflected in the decrease in the proportions of neutrophils (Fig. S5g-i) and monocytes (Fig. S5g-h, j) in spleen. These findings provide genetic evidence that fibroblast NNMT is essential for both initiation and reactivation of psoriatic inflammation.

**Figure 4.**
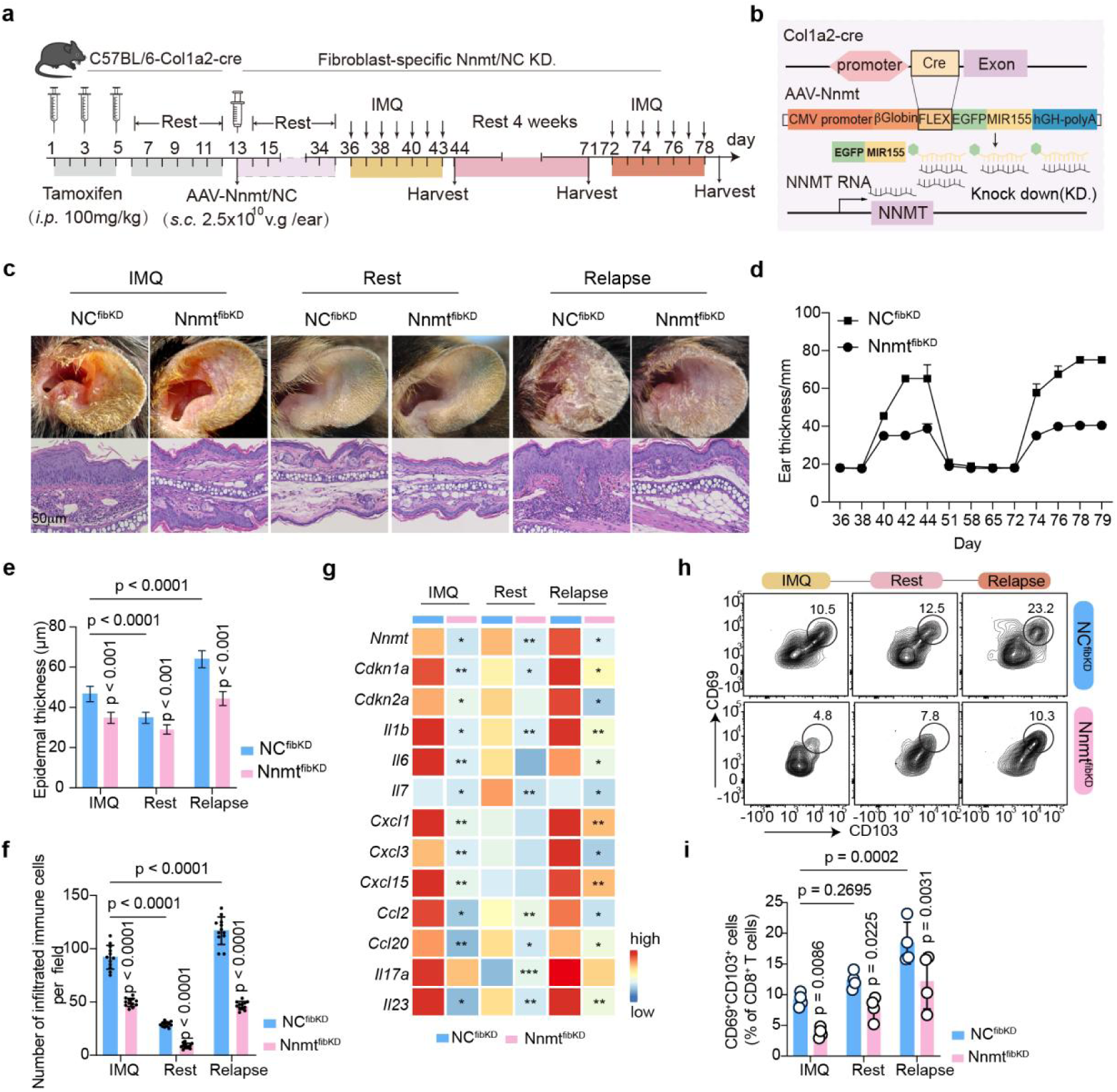
Fibroblast-specific NNMT deletion blocks IMQ-induced skin inflammation and relapse. **(a-b)** Schematics of generating fibroblast-specific Nnmt knockdown (Nnmt^fibKD^) strategy (a) and the vector sequence and the mechanism of AAV (b). **(c)** Representative appearance and H&E images and ear thickness of NC^fibKD^ and Nnmt^fibKD^ mice. **(d-f)** Statistical analysis of ear thickness (d), epidermal thickness (e) and infiltrated immune cells (f) (n= 12 of 4 skin samples/group). **(g)** Heatmap showing the expression of inflammatory genes detected by qRT-PCR from NC^fibKD^ and Nnmt^fibKD^ mice during the psoriasis phase, rest phase, and relapse phase (n= 4 skin samples/group). Data are presented as means and each group was compared with corresponding Nnmt^fibKD^ group in the same phase. **(h-i)** Representative images (h) and statistical analysis (i) of flow cytometry showing the proportion of skin Trm cells (CD103^+^ cells of CD8^+^ T cells) from NC^fibKD^ and Nnmt^fibKD^ mice during three phases (IMQ, Rest, and Relapse). The labeled p values above the bars represent the comparison with its corresponding control group in the same phase (n= 4 skin samples/group). *i.p.*, intraperitoneal injection. *s.c*., subcutaneous injection. v.g, viral genomes per milliliter. KD, knocked down. Data are represented as mean ± SD. Data in (e-g) and (i) were analyzed using two-way ANOVA. *p< 0.05, **p< 0.01.

### NNMT-SAM-H3K27me3 axis imprints fibroblast inflammatory memory

Given that NNMT catalyzes the methylation of nicotinamide using SAM as a donor (Fig. 5a), we hypothesized that NNMT-induced SAM depletion underlies the establishment of fibroblast memory. Indeed, enzyme-linked immunosorbent assay (ELISA) showed reduced SAM levels in NNMT-overexpressing fibroblasts (Fig. 5b). ATAC-seq of NNMT-overexpressing fibroblasts revealed increased genome-wide chromatin accessibility, particularly at promoter regions (47.72% of gained peaks) (Fig. S6a). Peak distribution (Fig. S6b) and heatmap (Fig. 5c) analyses also revealed increased accessibility within ± 1 kb of transcription start sites (TSS). Integration of ATAC-seq and RNA-seq datasets identified 85 genes exhibiting both increased promoter accessibility and transcriptional upregulation in NNMT-overexpressing fibroblasts, mainly enriched for SASP-associated cytokines such as *IL1B*, *IL6*, *IL11*, *CXCL3*, *CXCL8*, and *CCL7* (Fig. 5d). GO analysis revealed enrichment in pathways linked to cellular senescence, chemokine signaling, and leukocyte recruitment (Fig. 5e), indicating that NNMT drives fibroblast SASP by regulating inflammatory gene chromatin accessibility and subsequent transcription.

**Figure 5.**
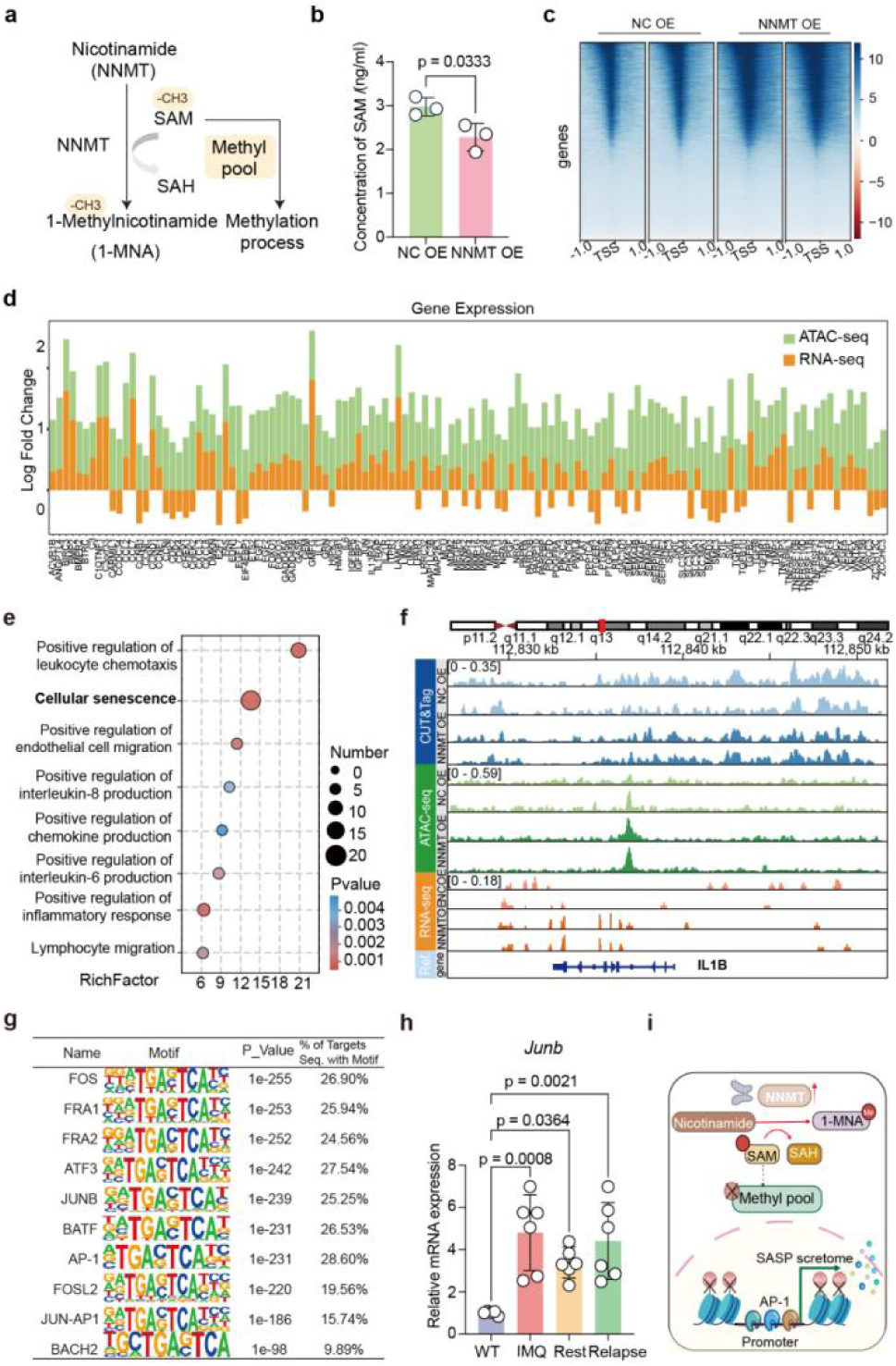
NNMT–SAM–H3K27me3 axis imprints fibroblast inflammatory memory. **(a)** Schematic of NNMT-catalyzed methyl-transfer using SAM as substrate. **(b)** ELISA of SAM levels in supernatants from NC/NNMT-OE fibroblasts (n= 3 cell samples/group). **(c)** Heatmap of TSS-centered signals (n= 2 cell samples/group). **(d)** Bar plot of intersection genes altered in ATAC-seq and RNA-seq (RNA-seq threshold: 1.2; ATAC-seq threshold: 1.5). **(e)** GO enrichment analysis of upregulated intersection genes; bubble size indicates gene count, and color indicates p-value. **(f)** Integrated CUT&Tag, ATAC-seq, and RNA-seq tracks for *IL1B*, showing chromatin accessibility and promoter/exon features. **(g)** HOMER Known Motif enrichment; motif logos show predicted base preferences and percentage of target sequences containing each motif. **(h)** QRT-PCR of *Junb* mRNA in mouse dermis across WT, IMQ, rest, and relapse phases (n= 6 from 3 skin samples/group). **(i)** Schematic model: NNMT depletes intracellular SAM and reduces H 3K 27me3 at inflammatory gene promoters, increasing chromatin accessibility. Upon psoriasis-related cytokine stimulation, stress-related and AP-1 transcription factors (*e.g.*, JunB) bind and drive inflammatory-gene expression and fibroblast SASP. TSS, transcription start site. The region within 1 kilobase (kb) from the TSS is considered the promoter region. HOMER, Hypergeometric Optimization of Motif EnRichment. Data are represented as mean ± SD. Data in (b) were analyzed using an unpaired two-tailed Student’s t-test. Analysis of data in (h) was performed using one-way ANOVA with Tukey’s post hoc test.

To investigate the epigenetic basis of this activation, CUT&Tag profiling demonstrated a significant reduction in H 3K 27me3 occupancy at these SASP gene promoters in NNMT-overexpressing fibroblasts (Fig. 5F and Fig. S6c-d), which was validated by western blotting (Fig. S5e). Motif analysis identified AP-1 family members (*e.g.* JunB, Fos and ATF 3) as dominant transcription factors enriched in these accessible regions (Fig. 5g). QRT-PCR of mouse dermis confirmed sustained *Junb* expression during both remission and relapse phases (Fig. 5h and Fig. S6f), indicating persistent transcriptional “placeholding”. Using JASPAR motif prediction (Matrix ID: MA0490.3), JunB binding sites were identified in the promoters of seven canonical SASP genes, such as *IL1B, IL6, IL11, CXCL3, CXCL8, CCL2*, and *CCL7*, all with relative scores > 80% (Supplementary Table 4). Moreover, supplementation with exogenous SAM restored H 3K 27me3 (Fig. S5g) and suppressed expression of SASP-associated genes, including *IL1B*, *IL6*, *IL11*, and *CXCL8* (Fig. S6h-i). Thus, NNMT-driven SAM depletion leads to loss of repressive histone marks, increased chromatin accessibility, and sustained AP-1 binding, collectively imprinting fibroblast inflammatory memory (Fig. 5i).

### Pharmacologic NNMT inhibition or SAM supplementation erases fibroblast memory and prevents relapse

To test whether NNMT-mediated fibroblast memory is reversible, we performed pharmacologic inhibition experiments and metabolic rescue *in vivo*. In the IMQ mouse model, local administration of a selective NNMT inhibitor (NNMTi) markedly alleviated erythema, scaling, and ear thickness (Fig. 6a-b) and reduced *Il1b* and *Il6* expression in dermal tissue (Fig. 6c) during both active and remission phases.

**Figure 6.**
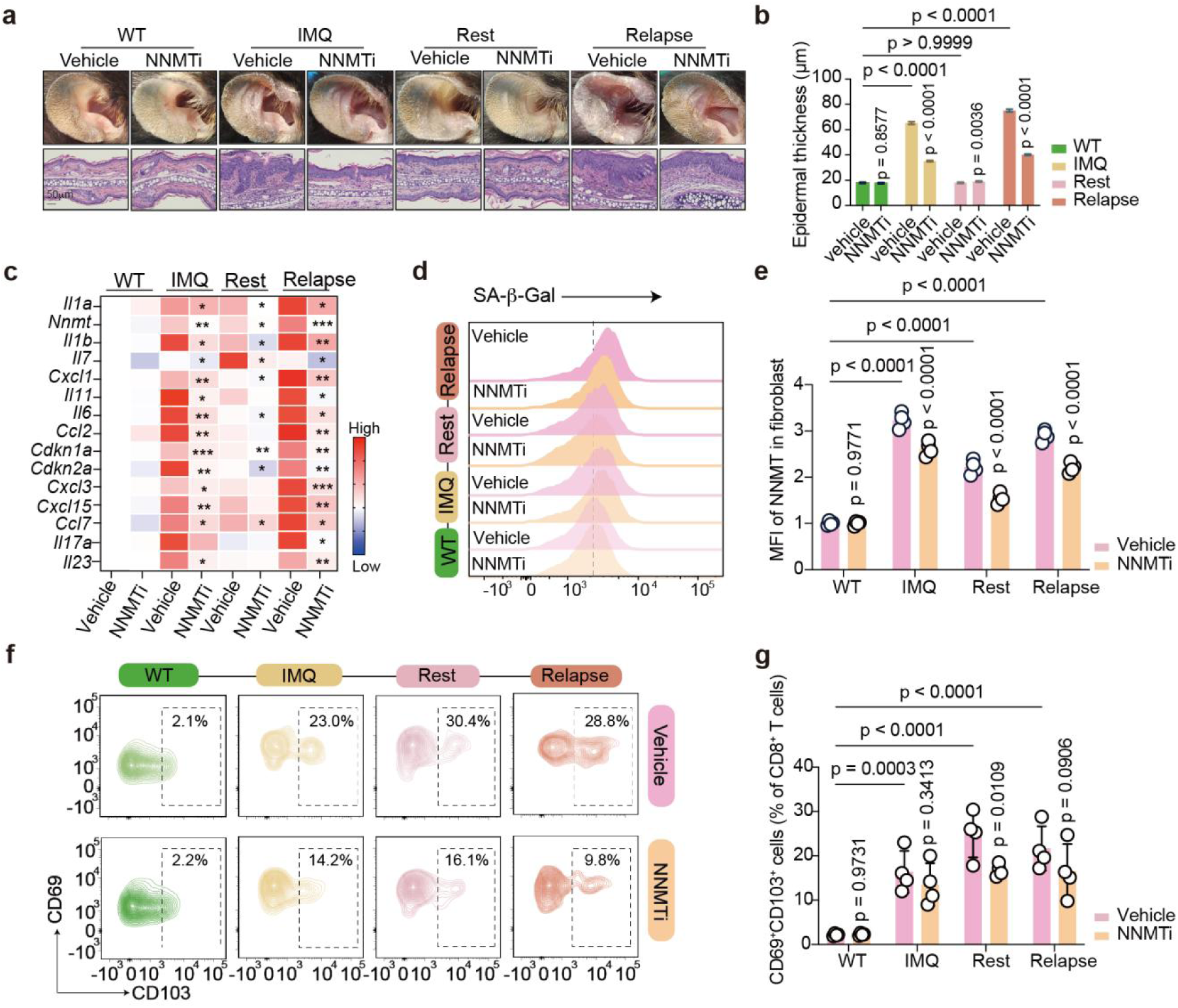
Targeting NNMT suppresses fibroblast SASP and prevents psoriasis relapse. **(a)** Representative appearance and H&E staining images. **(b)** Statistical analysis of epidermal thickness (n= 12 of 4 skin samples/group). **(c)** Heatmap of skin inflammatory cytokines expression detected by qRT-PCR in WT, IMQ, Rest, and Relapse groups (n= 4 skin samples/group). Data are presented as means. The mice treated with NNMT inhibitor are compared with their corresponding control in the same phase. **(d**-**e)** Representative images of flow cytometry (d) for mean fluorescence intensity of SA-β-Gal in fibroblasts and statistical analysis (e) (n= 4 skin samples/group). **(f-g)** Representative images (f) of flow cytometry and statistical analysis (g) of the proportion of CD8^+^CD103^+^ Trm of WT, IMQ, Rest, and Relapse mouse model with/without administration of NNMTi (n= 4 skin samples/group). Veh, vehicle. The labeled p values above the bars represent the comparison with its corresponding control group in the same phase. Data are represented as mean ± SD. Analysis of data was performed using two-way ANOVA. *p< 0.05, **p< 0.01, ***p< 0.001.

Flow cytometry revealed decreased proportions of SA-β-Gal⁺ fibroblasts (Fig. 6d-e) and CD 8⁺CD 103⁺ Trm cells in NNMTi-treated mice compared with controls (Fig. 6f-g). NNMTi-treated mice remained resistant to relapse following IMQ re-challenge. Similarly, daily intradermal SAM injections led to near-complete suppression of psoriasiform inflammation (Fig. S7a-c), reduced SASP gene expression (Fig. S7d), decreased the proportion of SA-β-Gal⁺ fibroblasts (Fig. S7e-f), and limited Trm infiltration (Fig. S7g-h). Collectively, these findings demonstrate that fibroblast inflammatory memory is metabolically and epigenetically reversible. In summary, targeting NNMT, either by restoring SAM availability or pharmacologic inhibition, can erase stromal memory, suppress Trm persistence, and achieve durable remission.

## Discussion

Many inflammatory skin disorders, including psoriasis, exhibit high rates of relapse at the same anatomical sites after withdrawal of targeted biologic therapies, underscoring the persistence of inflammatory memory that lies beyond the reach of current immune-directed treatments. Although CD 8⁺CD 103⁺ Trm cells are known mediators of localized disease recurrence^5, 6^, the mechanisms that sustain these cells within clinically resolved tissue microenvironment have remained incompletely defined. Here, we demonstrate that dermal fibroblasts acquire a persistent SASP during inflammation that reshapes the stromal microenvironment to promote CD 8^+^CD 103^+^ Trm differentiation and survival. Importantly, these senescent fibroblasts retain inflammatory memory through durable epigenetic remodeling. We identify NNMT as a key driver of this stromal memory program: by depleting SAM, NNMT reduces H 3K 27m3 deposition at SASP promoters and maintains chromatin accessibility, enabling sustained transcription factor engagement, including persistent JunB occupancy. Even after cytokine-driven senescence cues have been cleared during biologic therapy or remission, this epigenetically primed chromatin state lowers the threshold for disease reactivation. These findings challenge the prevailing immune-centric view of relapse and reveal that structural cells act as long-lived reservoir of pathological memory that actively instructs the immune niche.

The identification of a persistent stromal memory compartment represents a substantial conceptual advance. Although, psoriasis relapse has been associated with a “molecular scar”, in which inflammatory genes remain aberrantly expressed despite clinical resolution, memory in non-immune cells has been attributed to epigenetic alterations that prime rapid transcriptional responses. Most of these chromatin changes are transient, yet certain histone modifications, such as H 3K 4me1 at enhancers and H 3K 4me3 at promoters, can persist and maintain a poised regulatory landscape for future activation^16, 17^. Previous work has focused primarily on immune cell persistence and keratinocyte-intrinsic memory mechanisms^14^. In contrast, our single-cell and spatial transcriptomic analyses across the disease continuum reveal that fibroblasts represent the dominant stromal cells, acquiring a stable SASP. Notably, this SASP program persists in fibroblasts isolated from clinically resolved skin after prolonged IL-17A blockade, indicating a fibroblast-intrinsic memory state that is not eliminated by current cytokine-targeted therapies. Functional studies in our murine model of psoriasiform relapse further support this conclusion. Thus, during remission fibroblasts retain elevated SA-β-Gal activity and continue to express pro-inflammatory mediators, including *Il6*, *Il7*, and *Il1b*. The spatial confinement of this SASP program to anatomically recalcitrant sites provides a mechanistic explanation for the clinical pattern of site-specific recurrence.

We further delineate the functional consequences of this stromal memory in establishing a pro-relapse niche. Although immune-stromal interactions have been described in inflammatory skin diseases, the hierarchical relationship and instructional capacity of stromal cells have remained unclear. Our computational deconstruction analyses position senescent fibroblasts as central signaling hubs within the lesional microenvironment. Functionally, we demonstrate that these fibroblasts exert a direct instructional influence on T cell fate, with conditioned media from SASP fibroblasts sufficient to induce CD 103 expression on CD 8⁺ T cells, thereby promoting Trm differentiation. These findings establish fibroblasts as active architects of the immune microenvironment that sustain Trm populations implicated in disease relapse.

Central to this stromal memory program is NNMT, which we identified as the most significantly upregulated gene in senescent fibroblasts. NNMT functions as a metabolic rheostat by consuming SAM thus manipulating the intracellular “methylation sink”, and thereby reshaping histone methylation landscapes. Its role as a master metabolic regulator of fibroblast pathogenicity is increasingly appreciated across tissues - in cancer-associated fibroblasts, NNMT drives epigenetically encoded immunosuppressive programs that recruit myeloid-derived suppressor cells and impair CD 8⁺ T cell effector function^18, 19^. Analogously, our data establish NNMT as the central regulator of the SASP program in psoriatic fibroblasts. Both gain- and loss-of-function studies demonstrate that NNMT is necessary and sufficient to induce the full SASP phenotype and to endow fibroblasts with the capacity to instruct Trm-like cell differentiation. These findings position NNMT as a pivotal molecular node linking fibroblast-intrinsic epigenetic remodeling to the establishment of a pro-relapse immune niche.

At a mechanistic level, we define the NNMT-SAM-H 3K 27me3 axis as the central epigenetic driver for fibroblast-mediated inflammatory memory. Senescent cells undergo extensive chromatin reorganization^12, 20^, and the heterochromatin mark H 3K 27me3 is a key pressive modification that enforces transcriptional silencing. Its erosion, during senescence, has been implicated in the activation of pro-inflammatory programs^12,21^. EZH 2, the methyltransferase responsible for depositing H 3K 27me3, is known to restrain SASP gene expression^22^. Consistent with this framework, we found that NNMT-dependent depletion of SAM diminishes H 3K 27me3 at SASP loci, thereby releasing repression of IL6, IL1B, and CXCL8, and other inflammatory mediators. This mechanism parallels observations in aging muscle stem cells, where SAM loss reduces heterochromatin integrity and supplementation restores function^23^.

Our dissection of the fibroblast SASP reveals nuanced transcriptional regulation among individual components, providing deeper insight into the maintenance of the pro-relapse niche. Intriguingly, while IL-7 was not among the most robustly elevated cytokines during active inflammation in our *in vitro* SASP model or in the acute phase of IMQ-induced dermatitis *in vivo,* which is similar to the findings conducted by Liu *et al*^8^. Its expression in dermal fibroblasts demonstrated a unique pattern of persistence, failing to resolve to baseline levels during the remission phase. This specific persistence of IL-7, a cytokine with a well-established non-redundant role in promoting the survival and maintenance of CD 8⁺ T cells, finds a compelling functional correlate in the biology of pathogenic Trm17 cells. As highlighted by recent work, the differentiation and tissue residency of CD 8⁺ Trm17 cells are strictly IL-7-dependent, orchestrated through the ICOS-c-Maf-IL-7 axis^5^. Recent work has solidified the role of IL-7 in psoriatic memory; for instance, Zareie *et al*. demonstrated that CD 8 ⁺ CD 103 ⁺ Trm cells in resolved psoriatic lesions are maintained by local survival signals, with IL-7R signaling being critically involved^24^. Our data align with and extend this model by proposing that the NNMT-stabilized epigenetic landscape in fibroblasts selectively maintains a permissive state for IL-7 transcription, among other factors, even after the resolution of overt inflammation. This creates a low-level but durable survival signal for the previously established Trm pool, effectively “locking in” the cellular mediators of recurrence. Thus, the persistence of IL-7 exemplifies how the NNMT-driven SASP program encompasses not only factors that drive initial Trm differentiation but also those that ensure their long-term tenancy within the resolved skin, thereby cementing the stromal foundation for disease relapse.

Chromatin accessibility further revealed strong AP-1 motif enrichment within persistently open regions, accompanied by sustained *Junb* expression *in vivo*. These data support a two-step memory model in which early cytokine signaling engages pioneer factors such as STATs to initiate chromatin opening^25^, followed by AP-1 family members that maintain transcriptional activity and preserve accessibility^14^. In this model, persistent AP-1 occupancy serves as a molecular “placeholder” stabilizing a fibroblast-intrinsic memory state even after inflammatory cues have resolved. In addition to its epigenetic role, NNMT also consumes nicotinamide^26^, potentially linking NAD^+^ metabolism with mitochondrial function and SASP regulation. Although NAD^+^ depletion contributes to senescence^27^, its contribution to NNMT-driven memory remains to be fully established.

The translational implications of these findings are substantial. Fibroblast-specific deletion of NNMT completely prevented relapse *in vivo*, and both pharmacologic NNMT inhibition and SAM supplementation restored epigenetic homeostasis and achieved durable remission. In contrast to immune-targeted biologics, which suppress inflammatory signals without erasing underlying stromal memory, targeting NNMT directly dismantles the fibroblast-intrinsic epigenetic program that sustains relapse potential. These results highlight a therapeutic paradigm in which NNMT inhibition, along with or combined with IL-17/IL-23 blockade, may simultaneously extinguish immune activation and eliminate the stromal reservoir of inflammatory memory, offering the possibility of fundamentally extended remission.

In summary, our findings unify metabolic, epigenetic, and immunological mechanisms into a cohesive model in which NNMT-driven fibroblast memory serves as a fundamental determinant of disease recurrence. By linking SAM depletion to chromatin deregulation, persistent SASP activity, and the maintenance of pro-relpase niche, this work positions stromal memory, and not immune cells alone, as a central driver of psoriasis relapse. The widespread induction of NNMT across diverse chronic inflammatory and fibrotic conditions suggests that this pathway constitutes a generalizable stromal program. As such, targeting NNMT may offer a broadly applicable strategy to extinguish inflammatory memory and achieve durable remission in psoriasis and potentially other immune-mediated diseases.

## Code availability

This study did not generate a new algorithm or code. The custom code and scripts used for analyses and generating related graphs are available upon request.

## Author contributions

Conceptualization: Yifan Zhou, Yuming Xie, Kang Li, Chencan Su. Methodology: Yang Yang, Pei Qiao, Bingyu Pang, Yixin Luo, Jiaoling Chen, Hui Fang, Wanting Liu, Zhiguo Li, Yaxing Bai. Investigation: Yifan Zhou, Yuming Xie, Kang Li, Chencan Su. Visualization: Yifan Zhou, Yuming Xie, Kang Li, Chencan Su. Funding acquisition: Gang Wang, Shuai Shao. Project administration: Gang Wang, Shuai Shao, Johann E. Gudjonsson. Supervision: Gang Wang, Shuai Shao. Writing – original draft: Yifan Zhou. Writing – review & editing: Shuai Shao, Johann E. Gudjonsson.

## Additional information

The Supplementary Materials provided as PDF file includes Figs. S1 to S7 and Supplementary Methods (H&E staining, primary fibroblast culture and treatment, SA-β-Gal cell staining, immunofluorescence, isolation, activation, and induction of human CD 8^+^ T Cells, flow cytometry, RNA-seq library preparation, CUT&Tag library preparation, ATAC-seq library preparation, sequencing data processing, Enzyme-Linked Immunosorbent Assay (ELISA), RNA extraction and qRT-PCR analysis, and western blot Analysis). Supplementary Table 1-4 are provided as Excel file.

**Correspondence** and requests for materials should be addressed to Shuai Shao.

## Methods

### Research compliances

All analyses of human materials were done in full agreement with our institutional guidelines, with the approval of the Ethical committee of the Xijing Hospital, the Fourth Military Medical University (KY20222296-F-1). Written informed consent was obtained from each participant. This study is compliant with the Guidance of the Ministry of Science and Technology (MOST) for the Review and Approval of Human Genetic Resources. All animal procedures complied with the National Institutes of Health Guide for the Care and Use of Laboratory Animals and with Institutional Animal Care and Use Committee approval at the Fourth Military Medical University. Animal Experimental Ethical Inspection was approved Laboratory Animal Welfare and Ethics Committee of Fourth Military Medical University.

### Human skin sample

Human skin samples (punch biopsies) were obtained from healthy individuals and psoriasis patients at Xijing Hospital, Fourth Military Medical University (China). All patients enrolled in our study had no other autoimmune or systemic diseases. Patients were not on any systemic treatment for a month and had not used any biologics. All skin lesions/tissues and the surrounding 5 cm^2^ area were not treated with any therapeutic measures for at least 2 weeks before biopsy. Two dermatologists independently determined that the dissected skin tissues were representative of their disease type. Human blood samples were collected from healthy donors, outpatients, and inpatients, and were eligible to participate if they had plaque psoriasis for the first time, without other autoimmune or systemic diseases, and were not receiving systemic treatment or biologics for at least one month before blood sample collection.

### Mouse models

C57BL/6 J mice (6 – 8 weeks old) were purchased from Department of Laboratory Animal Medicine of the Fourth Military Medical University (Xian, Shaanxi, China). Both male and female were used for all experiments. Mice were randomly assigned to groups of 4 mice, then bred and maintained in specific pathogen-free conditions at a constant temperature and air humidity, under a standard 12-h light-dark paradigm, with free access to food and water. For *in vivo* experiments, researchers were blinded to the treatment of each animal received until data were analyzed. After the experiments, mice were euthanized by an overdose of sodium pentobarbital.

To induce skin inflammation, the imiquimod (IMQ)-induced psoriasis-like mouse model was used. Mice were treated daily with topical applications of 2mg IMQ cream (5%IMQ, INova Pharmaceuticals, 3 m Health Care) on the ears for a consecutive 7 days. The epidermal thickness was measured in a blinded way after completion of the 7 days of IMQ treatment. By applying 2mg IMQ cream for 7 days, natural relief for 4 weeks, and applying 2mg IMQ cream again for 7 days, a mouse model of psoriatic inflammation relapse was established.

Fibroblast-specific NNMT knockdown mice using adeno-associated virus (AAV)-mediated delivery of a LoxP-flanked NNMT construct in B6.Cg-Tg(*Col1a2*-Cre^ERT^,-ALPP)7Cpd/J mice, purchased from Shanghai Model Organisms Center. To activate Cre recombinase, tamoxifen (Beyotime) was dissolved in corn oil at a concentration of 20 mg/mL. Mice received intraperitoneal injections of tamoxifen at a dose of 100 mg/kg body weight every other day for a total of three administrations. Following a one-week recovery period, fibroblasts in the skin were targeted via subcutaneous injection of a viral suspension into the ear. Based on the predetermined viral titer, a single injection of 40 µL of the viral solution, containing 2.5× 10^10^ viral genomes (*v.g.*) per ear, was administered. Three weeks post-injection, skin lesions from the injection site were harvested and processed for sectioning. Successful transduction of fibroblasts and subsequent Cre-mediated recombination was confirmed by visualizing green fluorescent protein (GFP) expression under a fluorescence microscope.

For local drug administration, mice were anesthetized daily by intraperitoneal injection of 1.5% weight/volume (w/v) tribromoethanol (0.02 mL/g body weight) and received a 50 µL subcutaneous injection into the ear pinna once daily for 7 consecutive days (both in IMQ and relapse phases), commencing 2 hours prior to the daily application of IMQ. To evaluate the effects of S-adenosylmethionine (SAM) (HY-B0617A, MCE), a stock solution (100 mg/mL in sterile saline) was prepared by sonication, sterile-filtered, and diluted to working concentrations of 0.4, 2, and 20 mg/mL for the 1, 5, and 50 mg/kg dose groups, respectively; control mice received the vehicle (sterile saline) alone. For NNMT inhibition studies, NNMT inhibitor (JBSNF-000088, MCE) was dissolved in a vehicle containing 5% dimethyl sulfoxide (DMSO) and 95% sulfobutyl-β-cyclodextrin (SBE-β-CD, 30% w/v in phosphate-buffered saline) to achieve a final concentration of 25 µg/mL, delivering 1.25 µg per ear. Control mice for this experiment received injections of the corresponding vehicle (5% DMSO in 95% SBE-β-CD solution) without the inhibitor.

### Single-cell RNA-seq analysis

Publicly available single-cell RNA-sequencing (scRNA-seq) datasets were downloaded from the Gene Expression Omnibus (GEO). All datasets were processed following a consistent analytical workflow to enable comparison of transcriptional alterations and cellular senescence levels under different conditions.

For each dataset, the expression matrices were imported into Seurat (version 5.1.0) and subjected to a unified analysis pipeline, including quality control, data normalization, identification of highly variable genes, dimensionality reduction, batch-effect correction, clustering, and cell-type annotation. For visualization of the GSE173706 dataset, dimensionality reduction was performed using the RunUMAP function, and annotated cell clusters were displayed using DimPlot.

Cellular senescence at the single-cell level was quantified using both the AUCell package (version 1.26.0) and the AddModuleScore function in Seurat based on the SenMayo gene set (Dominik Saul *et al*.), which has been widely used for senescence scoring. Fibroblasts were ranked by Cellular Senescence Score (CS score) and divided into a high-senescence group (top 40%) and a low-senescence group (bottom 40%). Differentially expressed genes (DEGs) between CS-high and CS-low fibroblasts were identified using the FindMarkers function with thresholds of |log2FoldChange| > 0.25 and adjusted p value < 0.05.

To investigate the dynamic progression of senescence, pseudo-time trajectory analysis was conducted using monocle2 (version 3.1.7). Trajectories were constructed based on DEGs between the CS-high and CS-low groups. Gene expression changes along developmental branches were visualized using the plot_genes_branched_pseudotime function.

For intercellular communication analysis between fibroblasts with distinct senescence states and T-cell subsets, T cells were re-annotated and classified into six subpopulations, including CD 103⁺ T cells. Cell–cell communication networks were inferred using CellChat (version 2.12), and interaction strengths were visualized using circle plots generated by the netVisual_circle function.

Transcriptomic datasets used in this study:

| Transcriptomic dataset | Accession numbers |
| --- | --- |
| Psoriasis | GSE173706 (including spatial transcriptomics), GSE198805, GSE121212, GSE225475 |
| Atopic dermatitis | GSE147424 |
| Lupus | GSE186476 |
| Scleroderma | GSE138669 |
| Keloid | GSE163973 |
| Normal scar | GSE163973 |
| Different body site | GSE202352, GSE130973, GSE202352 |

Antibodies
| Antibodies | Supplier | Catalog number | Dilution |
| --- | --- | --- | --- |
| rabbit-anti-P16<sup>INK4A</sup> antibody | Abcam | ab211542 | 1:100 |
| rabbit-anti-P21 antibody | Abcam | ab188224 | 1:100 |
| mouse-anti-Vimentin antibody | Abcam | ab8978 | 1:100 |
| rabbit-anti-NNMT antibody | Abcam | ab119758 | 1:100 |
| Goat Anti-Rabbit IgG H&L (Cy3) antibody | Abcam | ab97075 | 1:1000 |
| Goat Anti-Mouse IgG H&L (FITC) antibody | Abcam | ab97022 | 1:1000 |
| Goat Anti-Mouse IgG H&L (Cy3) antibody | Abcam | ab97035 | 1:1000 |
| Zombie UV<sup>TM</sup> Fixable Viability Kit | BioLegend | 423107 | 1:1000 |
| APC/Cyanine7 anti-mouse CD3ε antibody | BioLegend | 100329 | 1:100 |
| APC anti-mouse CD4 antibody | BioLegend | 100412 | 1:100 |
| PerCP/Cyanine5.5 anti-mouse CD8a antibody | BioLegend | 100734 | 1:100 |
| PE anti-mouse CD8a antibody | BioLegend | 162304 | 1:100 |
| PerCP/Cyanine5.5 anti-mouse CD19 antibody | BioLegend | 152406 | 1:100 |
| FITC anti-mouse CD44 antibody | BioLegend | 156008 | 1:100 |
| FITC anti-mouse CD45 antibody | BioLegend | 157214 | 1:100 |
| Pacific Blue anti-mouse CD69 antibody | BioLegend | 104524 | 1:100 |
| PE/Cyanine7 anti-mouse CD103 antibody | BioLegend | 121426 | 1:100 |
| PE/Cyanine7 anti-mouse CD140a antibody | BioLegend | 135912 | 1:100 |
| PE anti-mouse/human CD11b antibody | BioLegend | 101208 | 1:100 |
| FITC Anti-Mouse Ly6G antibody | BioLegend | RD21311F | 1:100 |
| PE/Cyanine7 Anti-Mouse Ly6C antibody | BioLegend | RD21514F | 1:100 |
| APC anti-human CD3 antibody | BioLegend | 300458 | 1:100 |
| FITC anti-human CD4 antibody | BioLegend | 357406 | 1:100 |
| PerCP/Cyanine5.5 anti-human CD8a antibody | BioLegend | 301032 | 1:100 |
| Pacific Blue anti-human CD69 antibody | BioLegend | 310920 | 1:100 |
| PE anti-human CD103 antibody | BioLegend | 350206 | 1:100 |
| Beta Tubulin Polyclonal antibody | Proteintech | 14555-1-AP | 1:500 |
| Histone H3 Antibody | Cell signaling technology | 9715 | 1:100 |
| H3K27me3 | Cell signaling technology | 9733 | 1:100 |

Hematoxylin-eosin staining (H&E) staining, primary fibroblast culture and treatment, SA-β-Gal cell staining, immunofluorescence, isolation, activation, and induction of human CD 8^+^ T Cells, flow cytometry, RNA-seq library preparation, CUT&Tag library preparation, ATAC-seq library preparation, sequencing data processing, Enzyme-Linked Immunosorbent Assay (ELISA), RNA Extraction and qRT-PCR Analysis, and western blot analysis are provided in Supplementary Materials.

### Reporting summary

Further information on research design is available in the Nature Portfolio Reporting Summary linked to this article.

### Statistics analysis

Each experiment was performed at least 3 times. All data are presented as the mean ± SD. Statistical analyses of the data were performed using GraphPad Prism (version 9.2.0) and R. The differences between the 2 groups were compared by an unpaired, two-tailed Student’s t-test. For comparisons of more than 2 groups, one-way ANOVA was performed. The differences among groups defined by two categorical factors were compared using two-way ANOVA. The number of samples and the specific analysis method are indicated in the legend. p< 0.05 was considered significant.

